# Topological confinement by a membrane anchor suppresses phase separation into protein aggregates: implications for prion diseases

**DOI:** 10.1101/2024.04.09.588656

**Authors:** Kalpshree Gogte, Simon Kriegler, Verian Bader, Janine Kamps, Prerna Grover, Roland Winter, Konstanze F. Winklhofer, Jörg Tatzelt

**Author notes:** To whom correspondence should be addressed: Jörg Tatzelt, Ruhr University Bochum, Universitätsstr. 150, D-44801 Bochum, Germany.

## Abstract

Protein misfolding and aggregation are a hallmark of various neurodegenerative disorders. However, the underlying mechanisms driving protein misfolding in the cellular context are incompletely understood. Here we show that the restriction of conformational degrees of freedom by a membrane anchor stabilizes the native protein conformation and suppresses liquid-liquid phase separation and protein aggregation. Inherited prion diseases in humans and neurodegeneration in transgenic mice are linked to the expression of anchorless prion protein (PrP), suggesting that the C-terminal glycosylphosphatidylinositol (GPI) anchor of native PrP impedes spontaneous formation of neurotoxic and infectious PrP species. Combining novel *in vitro* and *in vivo* approaches, we show that anchoring to membranes prevents spontaneous aggregation of PrP. Upon release from the membrane, PrP undergoes a rapid conformational transition to detergent-insoluble aggregates. Our study supports an essential role of the GPI anchor in preventing spontaneous misfolding of PrP^C^.

## INTRODUCTION

Toxic protein aggregation is a characteristic feature of various neurodegenerative disorders, such as Alzheimer’s, Parkinson’s, and prion diseases. Understanding the forces driving the conformational transition of natively folded proteins into pathogenic conformers is crucial for developing therapeutic strategies. Prion diseases in humans and other mammals are caused by a conformational transition of the cellular prion protein (PrP^C^) to a misfolded isoform called scrapie prion protein (PrP^Sc^), which is the main component of infectious prions (Prusiner, 1998). The spontaneous conversion of PrP^C^ into PrP^Sc^ is an extremely rare event, evidenced by an annual incidence of sporadic prion diseases in humans of only 1 to 2 cases per 1 million. In contrast, upon inoculation, even minuscule amounts of PrP^Sc^ efficiently causes prion diseases by interacting with endogenous host PrP^C^ and catalyzing its conversion to PrP^Sc^. Another etiology of prion diseases is associated with human prion protein gene (*PRNP*) mutations (Mead *et al*, 2019). Of particular interest is a nonsense mutation that does not affect the protein sequence of mature PrP^C^ but prevents its post-translational modification with a C-terminal glycosylphosphatidylinositol (GPI) anchor. As a result, full-length PrPΔGPI is secreted instead of being tethered to the outer leaflet of the plasma membrane. Strikingly, expression of PrPΔGPI cause inherited prion diseases in humans and spontaneous neurodegeneration in transgenic mice (Chesebro *et al*, 2005; Jansen *et al*, 2010; Stohr *et al*, 2011). These findings revealed that the membrane anchor suppresses the spontaneous conversion of PrP^C^ to PrP^Sc^. Notably, the core conformation of infectious amyloid formed by GPI-anchored PrP^Sc^ or ΔGPI-PrP^Sc^ does not substantially differ (Hoyt *et al*, 2022).

Many proteins associated with neurodegenerative diseases, such as α-synuclein, TDP-43, Tau and PrP, undergo liquid-liquid phase separation (LLPS) under physiological conditions, suggesting that aberrant phase transitions of proteins under pathophysiological conditions may be implicated in the formation of toxic amyloid (rev (Alberti & Dormann, 2019; Babinchak & Surewicz, 2020; Darling & Shorter, 2021; Zbinden *et al*, 2020)). LLPS and liquid-to-solid phase separation (LSPS) of PrP have been analyzed in several *in vitro* studies, however, a possible role of the C-terminal membrane anchor in the phase separation of PrP has not been addressed so far (Agarwal *et al*, 2021; Huang *et al*, 2020; Kamps *et al*, 2021; Kostylev *et al*, 2018; Matos *et al*, 2020; Passos *et al*, 2021; Polido *et al*, 2024; Ramos *et al*, 2023; Tange *et al*, 2021). Membrane anchoring can significantly affect phase separation of proteins. First, the charged headgroups of the membrane lipids may interact with proteins and modulate protein-protein and protein-solvent interactions. Second, confinement to a two-dimensional milieu can significantly lower the critical concentration at which LLPS occurs. Indeed, LLPS has been demonstrated for various membrane proteins and is apparently involved in the regulation of many physiological processes (rev in (Banjade & Rosen, 2014; Case *et al*, 2019; Ditlev, 2021; Snead & Gladfelter, 2019). Whether membrane attachment also affects aberrant phase transitions of aggregation-prone proteins associated with neurodegenerative diseases is not clear.

Here we addressed the role of a membrane anchor for the phase separation of PrP. Employing *in vitro* and *in vivo* models we show that a membrane anchor stabilized a soluble conformation of PrP and suppresses its conformational transition into aggregates. Aberrant phase separation into detergent-insoluble protein aggregates was induced by releasing PrP from the membrane or by an interaction of membrane-bound PrP with aggregated protein seeds.

## RESULTS AND DISCUSSIONS

### A membrane anchor suppresses liquid-solid phase separation *in vitro*

To address a potential role of the C-terminal GPI anchor in influencing phase separation of PrP, we established a new model system with membrane-anchored recombinant PrP. It is based on planar supported lipid bilayers (SLBs) composed primarily of the lipid POPC (1-palmitoyl-2-oleoyl-glycero-3-phosphocholine).The lipid DGS NTA-Ni (1,2-dioleoyl-sn-glycero-3-[(N-(5-am ino-1-carboxypentyl)iminodiacetic acid)succinyl], nickel salt) was included to anchor recombinant PrP via its carboxy-terminal hexa-histidine (6xHis) tag to the SLBs (Fig. 1A). To validate the fluidity of the outer layer of the SLBs we included 0.15mol% rhodamine DHPE (1,2-dihexadecanoyl-*sn*-glycero-3 phosphoethanolamine) and performed fluorescence recovery after photobleaching (FRAP) recordings (Fig. 1B). Recombinant PrP fusion proteins containing an N-terminal maltose-binding protein (MBP) to keep PrP soluble were used to study the phase separation. After purification, the N-terminal MBP was cleaved off by TEV protease to initiate the phase transition (Banik *et al*, 2024; Kamps *et al*., 2021; Polido *et al*., 2024; Ramos *et al*., 2023). For the current study, we additionally included a PreScission (3C) protease cleavage site, located between the GFP and the C-terminal His-tag, to allow the inducible release of PrP from the SLBs (Fig. 1A). In buffer with physiological salt concentration (10 mM Tris pH 7.4, 150 mM NaCl) PrP underwent phase separation and formed undynamic protein assemblies after release of the MBP by TEV protease (Fig. 2A, S1A), corroborating earlier results (Banik *et al*., 2024; Polido *et al*., 2024). We then anchored MBP-PrP-GFP via its C-terminal His-tag to the SLBs. After extensive washing to remove unbound MBP-PrP-GFP, we added TEV protease. In contrast to PrP-GFP in solution, membrane-bound PrP-GFP did not form assemblies after cleavage of MBP (Fig. 2B). To exclude the possibility that the absence of LLPS was due to impaired TEV protease-mediated processing, we analyzed the sample by Western blotting to detect the cleaved products (Fig. S1B).

**Figure 1.**
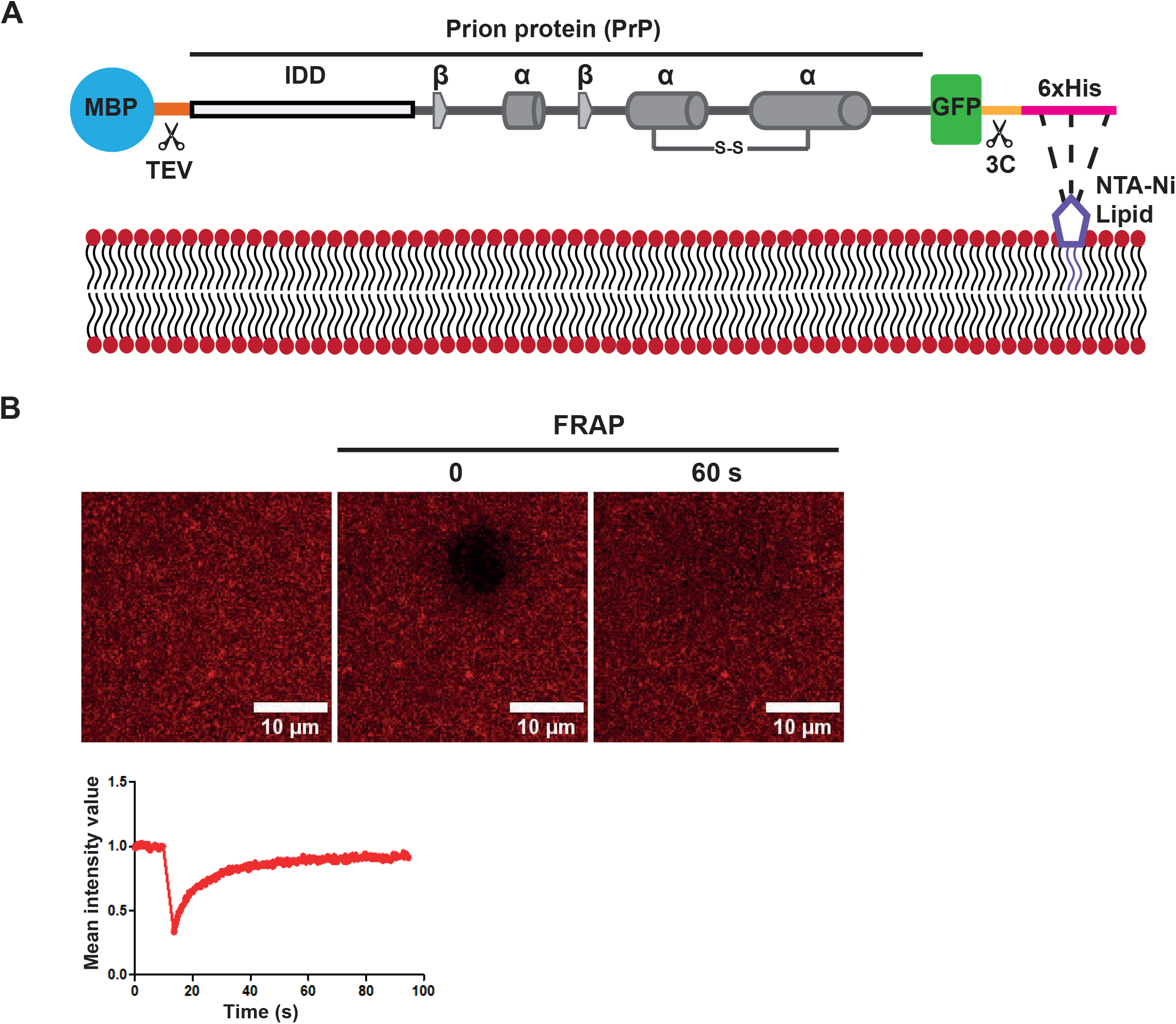
An *in vitro* model of membrane-anchored prion protein. **(A)** Schematic representation of the recombinant prion protein fusion protein (MBP-PrP-GFP) anchored to a supported lipid bilayer (SLB). The N-terminal maltose-binding protein (MBP) can be cut off with TEV protease. The C-terminal His-tag (6xHis) tethers the protein to the SLB via an interaction with NTA-Ni lipids. The protein can be liberated from the membrane via cleavage with 3C protease. IDD: intrinsically disordered domain; β: beta-strands; α: alpha-helices; S-S: disulfide bridge. **(B)** SLBs containing rhodamine DHPE were analyzed by laser scanning microscopy. Lipid mobility was measured by fluorescence recovery after photobleaching (FRAP). After 10 s of baseline recording (pre-bleach), a small area of interest (AOI) was photobleached. The average normalized fluorescence intensity of AOI was plotted over time.

**Figure 2.**
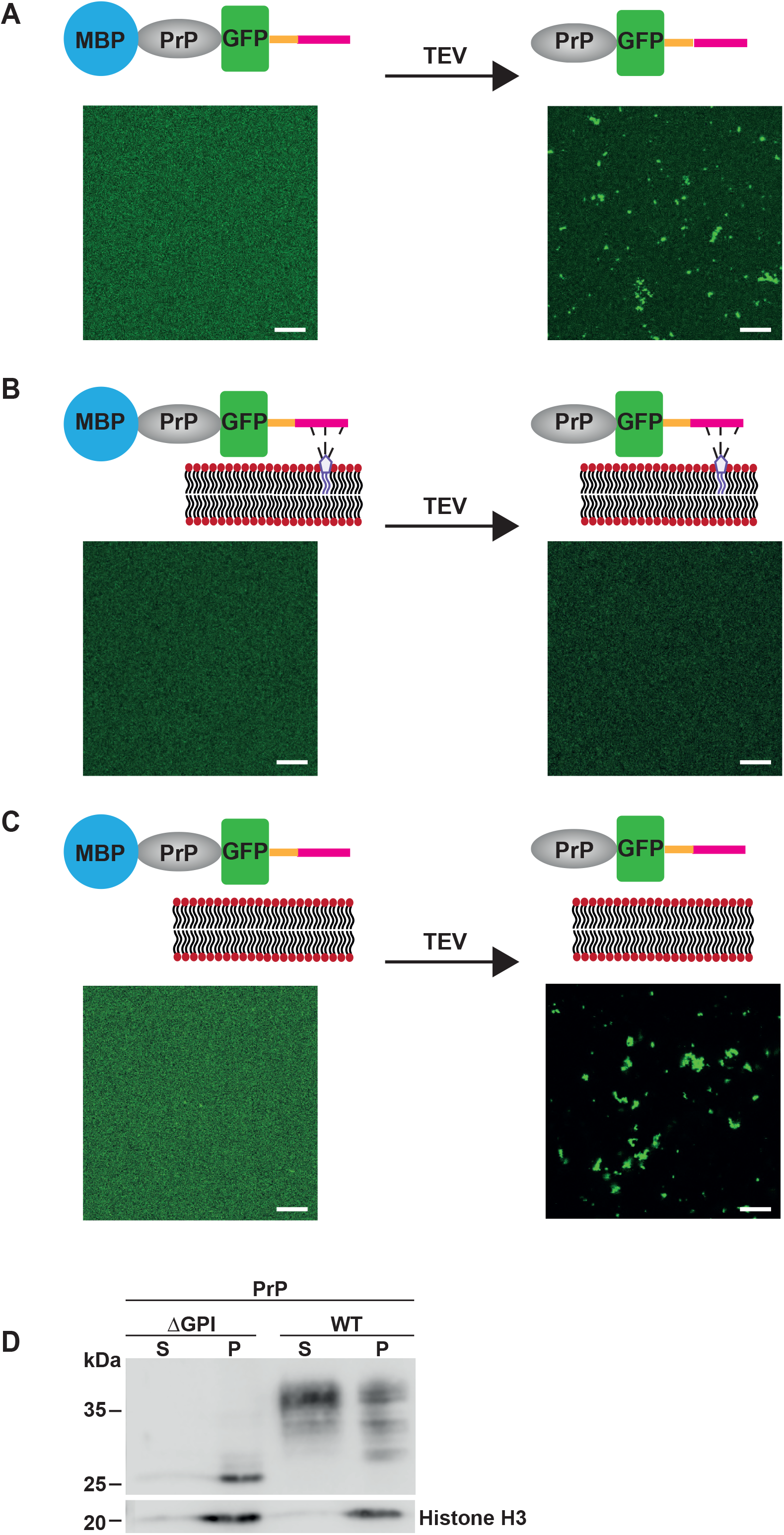
Membrane anchor prevents liquid-solid phase separation of PrP. **(A)** MBP-PrP-GFP (300 nM in 10 mM Tris pH 7.4, 150 mM NaCl) was incubated with TEV protease for 20 min and analyzed by laser scanning microscopy before and after TEV protease treatment (scale bar, 10 µm). **(B)** MBP-PrP-GFP (300 nM in 10 mM Tris pH 7.4, 150 mM NaCl) was tethered to NTA-Ni lipids in the SLB. After washing to remove unbound PrP, the SLB was incubated with TEV protease for 20 min and analyzed by laser scanning microscopy before and after TEV protease treatment (scale bar, 10 µm). **(C)** MBP-PrP-GFP (300 nM in 10 mM Tris pH 7.4, 150 mM NaCl) was added to SLBs lacking NTA-Ni. The sample was incubated with TEV protease for 20 min and analyzed by laser scanning microscopy before and after TEV protease treatment (scale bar, 10 µm). **(D)** N2a cells transiently expressing PrPΔGPI and wild-type (WT) PrP^C^ were lysed in detergent buffer and separated into a detergent-soluble (S) and -insoluble (P) fraction by centrifugation. The fractions were analyzed by Western blotting using an antibody against PrP and Histone H3 (loading control).

The question now was: Is the presence of the membrane interface sufficient to impede phase separation of PrP, or is the membrane anchor required? We therefore prepared SLBs without DGS NTA-Ni lipids to prevent the interaction of the C-terminal His-tag of PrP with the SLBs. We then added MBP-PrP-GFP and removed the MBP by TEV protease. Similar to PrP in pure solution, unanchored PrP rapidly aggregated in the presence of the lipid bilayer upon addition of TEV protease. Thus, strong membrane anchoring is required to suppress PrP aggregation (Fig. 2C).

Notably, the physiological relevance of these *in vitro* approaches is supported by previous *in vivo* studies. In cells, native GPI-anchored PrP^C^ is in a detergent-soluble conformation. In contrast, anchorless PrP (PrPΔGPI) spontaneously adopts a detergent-insoluble, partially protease K-resistant conformation (Blochberger *et al*, 1997; Kocisko *et al*, 1994; Rambold *et al*, 2006; Rogers *et al*, 1993; Walmsley *et al*, 2001; Winklhofer *et al*, 2003). This phenomenon is shown in Figure 2D. Transiently transfected neuronal cells expressing wild type PrP^C^ or PrPΔGPI were lysed in detergent buffer. After centrifugation the detergent-soluble and -insoluble fraction were analyzed by Western blotting. While the majority of GPI-anchored PrP^C^ partitioned into the detergent-soluble phase, PrPΔGPI adopted a detergent-insoluble conformation (Fig. 2D).

### Aggregation of membrane-bound PrP is induced by preformed seeds

In infectious prion diseases, preformed PrP^Sc^ interacts with GPI-anchored PrP^C^ and efficiently induces its conversion to PrP^Sc^. To mimic such a seeded aggregation of GPI-anchored PrP^C^ in our *in vitro* model, we tethered PrP to the SLBs, removed the MBP with TEV protease and then incubated the SLBs with aggregated recombinant PrP without a GFP tag (Fig. 3A). Indeed, the addition of preformed PrP aggregates rapidly induced aggregation of the membrane bound PrP-GFP (Fig. 3B). Moreover, the 3D reconstructions of the microscopic images revealed that the PrP-GFP assemblies were still attached to the SLBs (Fig. 3C).

**Figure 3.**
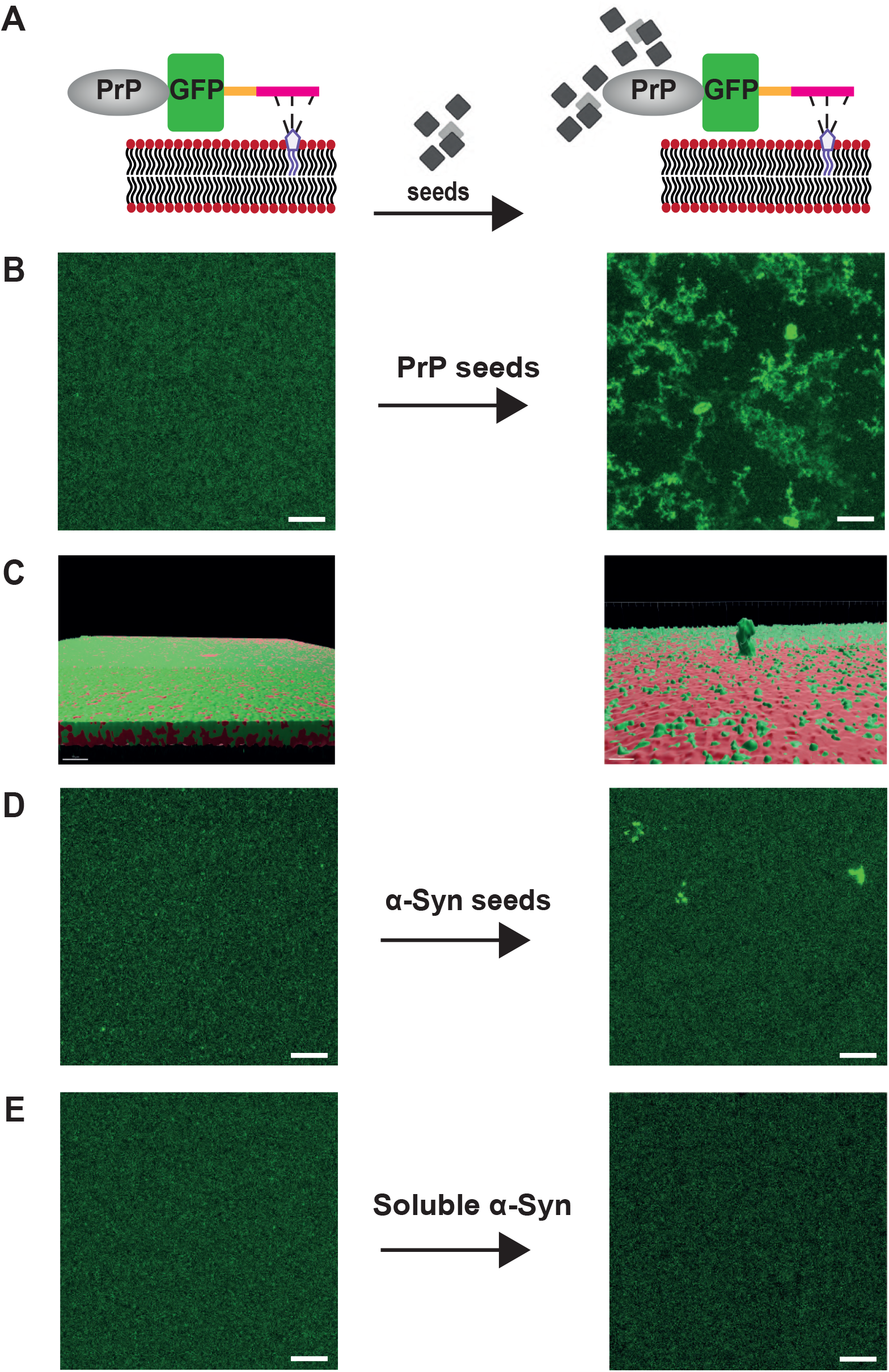
PrP and α-synuclein seeds induce aggregation of membrane-anchored PrP. **(A)** Schematic outline of the experiment. PrP was tethered to the SLBs and MBP was cleaved with TEV protease. The membrane bound PrP was then incubated with preformed seeds. **(B, C)** Membrane anchored PrP-GFP was incubated with preformed recombinant PrP seeds for 20 min. **(B)** The samples were analyzed by laser scanning microscopy before and after incubation with seeds (scale bar, 10 µm). **(C)** Volumetric 3D reconstructions generated with the IMARIS software. Red: SLBs, green: PrP-GFP. **(D)** Membrane anchored PrP-GFP was incubated with preformed recombinant α-Syn seeds for 20 min and analyzed by laser scanning microscopy before and after incubation with seeds (scale bar, 10 µm). **(E)** Membrane anchored PrP-GFP was incubated with soluble recombinant α-Syn for 20 min and analyzed by laser scanning microscopy before and after incubation (scale bar, 10 µm).

In addition to its role as a precursor for the formation and propagation of infectious prions, neuronal GPI-anchored PrP^C^ mediates neurotoxic effects of Scrapie prions (Brandner *et al*, 1996; Chesebro *et al*., 2005; Mallucci *et al*, 2003; Rambold *et al*, 2008), amyloid beta (Aβ) (Lauren *et al*, 2009; Nygaard & Strittmatter, 2009; Purro *et al*, 2018; Resenberger *et al*, 2011; Smith & Strittmatter, 2017), α-synuclein (α-Syn) (Ferreira *et al*, 2017), and Tau (Ondrejcak *et al*, 2018) via interaction of its N-terminal domain with β-sheet-rich oligomeric conformers of the respective misfolding-prone proteins. Whether such an interaction is associated with a conformational change of PrP^C^ has not yet been conclusively clarified (Konig *et al*, 2021; Kostylev *et al*., 2018). To address this question, we added preformed α-Syn aggregates to SLB-bound PrP-GFP. Similar to the PrP aggregates α-Syn aggregates induced aggregation of membrane-bound PrP, albeit with a lower efficiency (Fig. 3D). As a control, we show that soluble α-Syn did not induce aggregation of PrP (Fig. 3E). To our knowledge, these results are the first experimental evidence with purified components showing that pathogenic protein aggregates can induce misfolding of membrane-anchored PrP.

### Release from the membrane triggers PrP aggregation *in vitro* and *in cellulo*

To test the behavior of PrP after its release from the membrane, we made use of the PreScission (3C) protease cleavage site in front of the C-terminal His-tag. We first anchored MBP-PrP-GFP via its C-terminal His-tag to the NTA-Ni lipids of the SLBs and removed MBP with TEV protease-mediated cleavage. 3C protease was then added to cleave the C-terminal His-tag and to liberate PrP-GFP from the SLBs. Strikingly, PrP-GFP spontaneously formed aggregates after release from the membrane (Fig. 4A, S1C). 3D reconstructions illustrated that some of the PrP aggregates induced by addition of the 3C protease were not attached to the SLBs (Fig. S1D). When MBP was not removed prior to membrane release, the unanchored MBP-PrP-GFP did not form assemblies (Fig. S1E).

**Figure 4.**
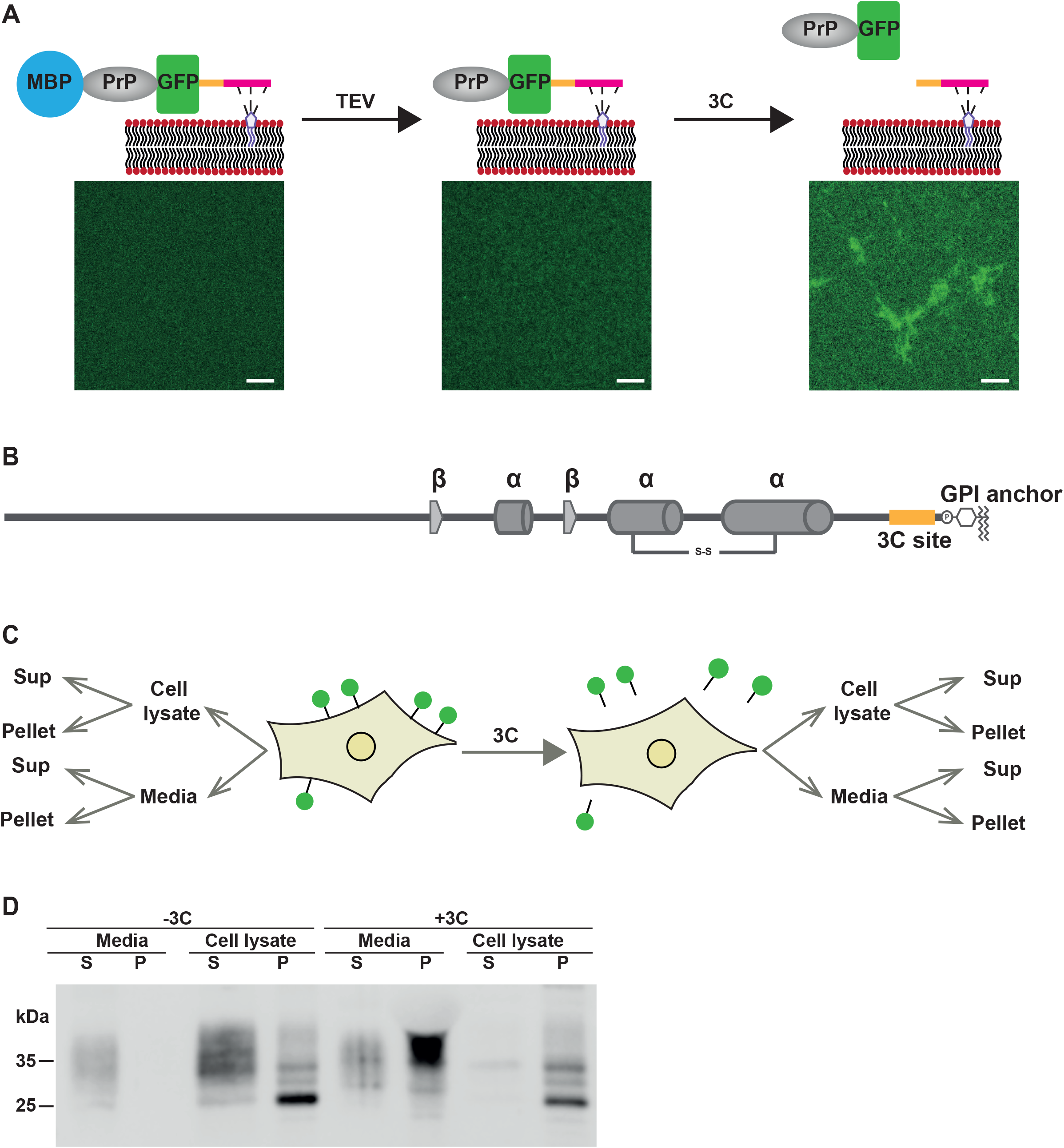
Membrane release induces the spontaneous formation of PrP aggregates. **(A)** Membrane anchored MBP-PrP-GFP (300 nM in 10 mM Tris pH 7.4, 150 mM NaCl) was first incubated with TEV protease to cleave MBP and then incubated with 3C protease to release PrP-GFP from the membrane. The samples were analyzed by laser scanning microscopy before and after the addition of TEV protease and after incubation with 3C protease (scale bar, 10 µm). **(B)** Scheme of GPI-anchored PrP with the 3C protease cleavage in front of the GPI anchor (PrP-3C-GPI) **(C)** Schematic outline of the experimental approach. N2a cells transiently expressing PrP-3C-GPI (shown in green on the cell surface) were incubated with 3C protease to release PrP-3C into the media. The cell lysates and the media were fractionated into detergent-soluble (Sup) and -insoluble fractions (Pellet) and analyzed by Western blotting. **(D)** N2a cells transiently expressing PrP-3C-GPI were treated with 3C protease (3 mg/ml) or PBS (-3C) for 4 h at 37 °C. Cell lysates and media were separated into a detergent - soluble (S) and -insoluble (P) fraction and analyzed by Western blotting using an antibody against PrP.

To corroborate our *in vitro* findings *in vivo*, we generated another construct for the inducible release of GPI-anchored PrP from the plasma membrane of mammalian cells. To achieve this, we introduced the 3C protease cleavage site in front of the C-terminal GPI anchor signal sequence (PrP-3C-GPI) (Fig. 4B). To test whether the release from the plasma membrane is accompanied by a conformational transition of PrP live N2a cells transiently expressing PrP-3C-GPI were cultivated for 4 h in the presence of 3C protease. To analyze the conformation of PrP, both cell lysates and the conditioned media were separated into detergent soluble and insoluble fractions and analyzed by Western blotting (Fig. 4C, D). In the absence of 3C protease, GPI-anchored PrP-3C was mainly found in the detergent-soluble fraction of the cell lysate, similar to wildtype PrP^C^ (Fig 2D) and only a small fraction of PrP-3C was detected in the conditioned media (Fig. 4D, -3C). In samples prepared from 3C-treated cells, PrP-3C was now present in the media, indicating that the 3C protease had liberated PrP-3C from the outer leaflet of the plasma membrane. Strikingly, a significant fraction of PrP-3C in the media was in the detergent-insoluble fraction, revealing a conformational transition of the membrane-bound detergent-soluble PrP into a detergent-insoluble conformation after its release from the membrane (Fig. 4D, +3C).

Our study demonstrated that a membrane anchor efficiently suppresses aggregation of PrP into aggregates. Spontaneous misfolding of PrP was triggered by releasing PrP from the membrane *in vitro* and in a cellular model. Moreover, misfolding of membrane-anchored PrP was observed after adding preformed protein aggregates as seeds (Fig. 5). These results provide a mechanistic basis for several interesting observations in prion biology. First, these findings help to understand why the expression of an anchor-less prion protein in humans and transgenic mice leads to the spontaneous formation of infectious prions. Second, our study also revealed that preformed protein aggregates induce a conformational conversion of membrane-anchored PrP. Such a seeded aggregation underlies the pathogenesis of infectious prion diseases. In addition, the conformational transition of GPI-anchored PrP upon engagement with extracellular protein aggregates may be associated with its pathophysiological role as a membrane receptor for toxic signaling of scrapie prions and other β-sheet-rich pathogenic protein assemblies (Ferreira *et al*., 2017; Lauren *et al*., 2009; Ondrejcak *et al*., 2018; Rambold *et al*., 2008; Resenberger *et al*., 2011 ).

**Figure 5.**
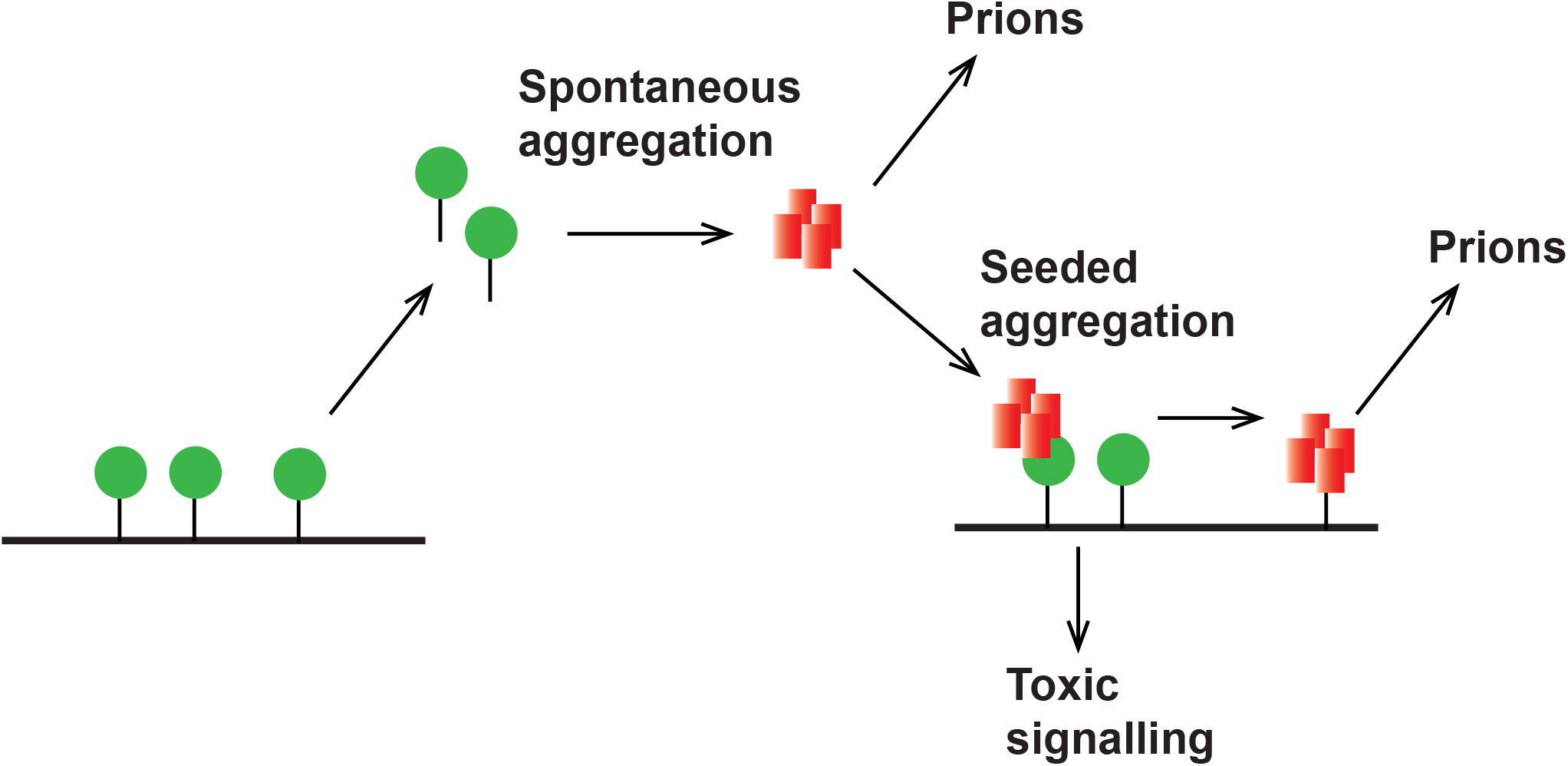
Schematic summary of the findings. The membrane anchor stabilizes the native conformation of PrP^C^ (green). After membrane release PrP spontaneously misfolds into aggregates that could interact with and induce misfolding of membrane-anchored PrP^C^. This may initiate formation of infectious prions or may initiate toxic signaling by the PrP^C^/Seed complex at the plasma membrane.

At first glance, our results may appear to contradict several compelling studies showing that membrane-anchored proteins efficiently undergo LLPS, even at much lower concentration than those at which LLPS occurs in pure solution. However, upon closer look it becomes apparent that the published studies are based on multi-component systems in which a membrane protein forms biomolecular condensates after an interaction with additional non-membrane-anchored proteins. Phase separation of membrane-anchored PrP is also seen in our system after addition of non-anchored preformed protein aggregates. Importantly, the concentration of membrane-bound PrP was sufficient to promote phase separation. This is illustrated by the rapid formation of PrP assemblies after release from the membrane.

How can the role of the membrane anchor in suppressing phase separation of PrP despite the high concentration of membrane-bound protein be explained? Most likely, the restricted conformational degrees of freedom of membrane-anchored PrP limit the potential for the formation of the multitude of intermolecular interactions that underly LLPS and are involved in protein aggregation. In addition, the conformation of the GPI-anchored PrP may be different to that of non-anchored PrP, i.e. the membrane-bound protein may adopt a different conformational substate. Indeed, a conformational change after membrane release was reported previously for a number of other GPI-anchored proteins (Barboni *et al*, 1995; Brewis *et al*, 1994; Durbin *et al*, 1994; Gmachl *et al*, 1993; Lehto & Sharom, 1998; Muller & Bandlow, 1994; Wang *et al*, 1996).

## MATERIAL AND METHODS

### DNA constructs

Plasmid maintenance and amplification was carried out using *Escherichia coli* TOP10^©^ (ThermoFisher Scientific). All PrP constructs were generated by standard PCR cloning techniques and are based on the coding region of mouse PRNP (GenBank accession number M18070) modified to express PrP-L108M/V111M, allowing detection by the monoclonal antibody 3F4. To generate MBP-TEV-PrP-GFP-3C-6His we exchanged the Tobacco etch virus (TEV) protease cleavage site between PrP-GFP and 6His with a PreScission protease (3C) site in plasmid MBP-TEV-PrP-GFP-TEV-6His used elsewhere (Kamps *et al*, 2021). The following DNA oligonucleotides were used with BsrgI and HindIII restriction sites.

Oligo with 3C: 5’ ATATGTACAAGCATCATCATCATCATCACTGAAAGCTTATA 3’

Oligo with 3C: 3’ TATAAGCTTTCAGTGATGATGATGATGATGCTTGTACATAT 5’

All the mammalian expression plasmids were cloned into pcDNA3.1(+)-Neo vector. PrPΔGPI: aa 1-226; PrP-3C: wild type PrP with the PreScission protease (3C) inserted after aa 223.

### Cell culture and transfection

Mouse neuroblastoma (N2a) cells were cultured in DMEM (1X) GlutaMAX™ supplemented with 10% (v/v) fetal bovine serum (FBS), 1% non-essential amino acid, 100 IU/ml penicillin and 100 μg/ml streptomycin sulphate under humidified conditions at 37 °C with 5 % CO_2_. For 3C protease treatment, cells were incubated in 3 mg/ml of 3C in 1X PBS for indicated time at 37 °C. Unless described otherwise, N2a cells were transfected using Lipofectamine 2000

(Invitrogen) according to the manufacturer’s instructions.

### Expression and purification of recombinant MBP-PrP fusion proteins

Plasmid maintenance, bacterial expression, and purification of MBP-PrP-GFP and MBP-PrP was performed as described earlier (Kamps *et al*., 2021). For protein expression 1 L transformed Origami™ B (DE3) cells in lysogeny broth (LB) medium was grown to an OD (600 nm) of 0.9 to 1.0. Expression was induced with 0.5 mM IPTG and incubated over night at 25 °C, 120 rpm. Bacteria were harvested by centrifugation (5,000g, 4 °C, 20 min), pellet was washed with 20 ml millipore water and centrifuged again (2,000g, 4 °C, 20 min). Pellets were stored at -20 °C until further use. For purification bacterial pellet was resuspended in lysis buffer (50 mM Na_2_HPO_4_/NaH_2_PO_4_ (pH 8.0), 500 mM NaCl, 0.01 mM ZnCl_2_, 10 % glycerol). Protein lysis was performed via SLM AMINCO French Press (Thermo Fisher Scientific, Waltham, MA) and the protein solution was centrifuged (40,000g, 45 min, 4 °C). The supernatant was loaded on a His-Trap FF column (GE Healthcare, Chicago, I L) equilibrated with lysis buffer and washed with 3 CV lysis buffer containing 20 mM imidazole and washed again with 3 CV lysis buffer containing 50 mM imidazole. Proteins were eluted with lysis buffer containing 200 mM imidazole and dialyzed over night at 4 °C in dialysis buffer (50 mM Na_2_HPO_4_/NaH_2_PO_4_ (pH 8.0), 500 mM NaCl, 0.01 mM ZnCl_2_, 5 % glycerol). The protein concentration was determined by NanoDrop 2000 (Thermo Scientific, USA), aliquoted and stored at -80 °C until further use.

### Generation of seeding-competent α-Syn and PrP aggregates

The generation of seeding-competent α-Syn aggregates have been described previously (Furthmann *et al*, 2023). In brief, an aliquot containing 1 ml α-Syn A53T with a concentration of 5 mg/ml (50 mM Tris, pH 7.5, 150 mM KCl) was thawed on ice and centrifuged (20,000g, 30 min, 4 °C). The supernatant was transferred into a new tube and incubated on a thermomixer for 24 h at 37 °C, 1000 rpm. The sample was divided into 20 µl aliquots and stored at −80 °C until further use. As described (Polido *et al*., 2024), PrP seeds were produced as follows. 10 μM MBP-PrP in aggregation buffer (10 mM Tris pH 7.4, 150 mM NaCl) was treated with TEV protease to cleave MBP and induce phase separation of PrP into aggregates.

### Preparation of small unilamellar vesicles (SUVs)

POPC (1-palmitoyl-2-oleoyl-*sn*-glycero-3-phosphocholine), DGS NTA-Ni (1,2-dioleoyl-sn-glycero-3-[(N-(5-am ino-1-carboxypentyl)iminodiacetic acid)succinyl], nickel salt), rhodamine DHPE (Invitrogen™ Lissamine™ Rhodamine B 1,2-dihexadecanoyl-*sn*-glycero-3 phosphoethanolamine, triethylammonium salt) stock solutions were prepared in chloroform and stored at 4 °C. To prepare SUVs, the components were mixed and lyophilized overnight. The components were dissolved in 10 mM HEPES pH 7.5 and sonicated for 20 min until the solution was clear. The mixture was passed through a membrane (PC Membranes 0.1μm, Avanti Polar Lipids) at least 21 times, using an extruder (Extruder Set with Holder/Heating Block, Avanti Polar Lipids) at 35 °C.

### Preparation of supported lipid bilayers (SLBs)

Coverslips were incubated in a glass chamber in H_2_O_2_ and H_2_SO_4_ (3:7 ratio) for 8 min. After extensive rinsing with millipore water, the coverslips were dried under N_2_ gas stream. Silicone gaskets (Press-to-seal™ silicone insulator with adhesive, ThermoFisher Scientific) punched with a 5-mm-diameter holes were stuck to the cleaned coverslips and left to bind for 15 min. 50 µl SUV solution was added to the wells for 20 min. Unbound vesicles were washed carefully with 1 ml of 10 mM HEPES pH 7.5, 150 mM NaCl. For SLBs functionalized with MBP-PrP-GFP, the protein was incubated with SLB for 20 min followed by a wash with 1 ml of 10 mM Tris pH 7.4, 150 mM NaCl to remove unbound protein.

### Microscopy

Fluorescent imaging laser scanning microscopy on a microscope (ELYRA PS.1; Carl Zeiss) with an imaging detector (LSM 880; Carl Zeiss) was performed. For z-stack scanning, a 63× numerical aperture 1.4 oil-immersion objective was used to record a stack of 134.95 × 134.95 × 8.39 μm/10.3 μm and 0.330/0.641 μm for each optical section. Depending on the presence or absence of assemblies, the total number of slices in the z-stack were modified. The argon laser power was set to 0.15 % at 488 nm and 0.30 % at 561 nm with a pixel dwell time of 0.38 μs.

### Fluorescence recovery after photo-bleaching (FRAP) recordings

For FRAP experiments, ZEN2.1 bleaching and region software module and a Plan-Apochromat 63× numerical aperture 1.4 oil differential interference contrast M27 objective was used. For regions of interest of SLB or PrP-GFP assembly, two circles of 52.35 μm^2^ or 8.05 μm^2^ area were chosen, respectively. One region was bleached with 100 % laser power and a pixel dwell time of 1.61 ms, with a scan time of 111.29 ms, and the other region was used as the reference signal. Data calculation was performed in Excel 2016, and diagrams were made with GraphPad Prism.

### Cell culture assays

#### Detergent solubility assay

As described earlier (Tatzelt *et al*, 1996) cells were washed twice with cold PBS, scraped off the plate, and pelleted by centrifugation. To separate detergent-soluble and -insoluble fractions the cells were lysed in cold detergent buffer (0.5% Triton X-100 and 0.5% sodium deoxycholate [DOC] in PBS) and centrifuged at 20,000g for 30 min at 4 °C. The supernatant (Sup) and pellet were analyzed by Western blotting. For the analysis of protein in the conditioned media, cells were treated with 3C protease (3 mg/ml) in 1 ml PBS. The media were collected and spun at 200g for 10 min at 4 °C to pellet down cell debris. After adjusting the media to 0.5% Triton/DOC, they were centrifuged at 20,000g for 30 min at 4 °C. Proteins present in the supernatant were TCA precipitated before the Western blot analysis.

### Immunoblotting

Proteins were fractionated by SDS-PAGE and transferred to nitrocellulose membranes by electroblotting. The nitrocellulose membranes were blocked with 5 % non-fat dry milk in TBST (TBS containing 0.1 % Tween 20) for 60 min at room temperature and subsequently incubated with the primary antibody diluted in blocking buffer for 16 h at 4 °C. After extensive washing with TBST, the membranes were incubated with horseradish peroxidase-conjugated secondary antibody for 60 min at room temperature. Following washing with TBST, the antigen was detected with the enhanced chemiluminescence (ECL) detection system (Promega) as specified by the manufacturer with Azure Sapphire Biomolecular Imager (Azure Biosystems, USA).

---

**Supplementary Figure 1.**
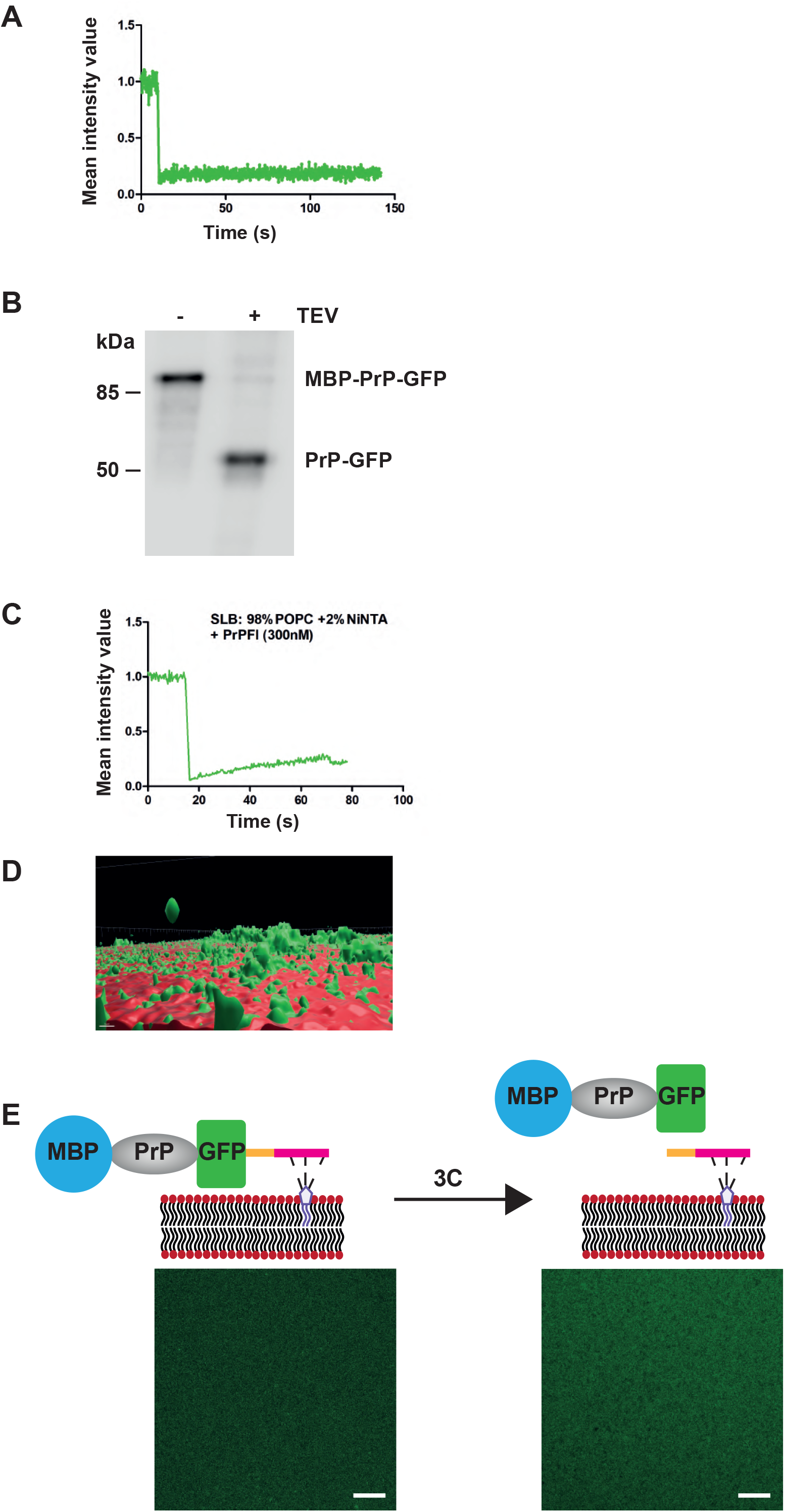
**(A)** MBP-PrP-GFP (300 nM in 10 mM Tris pH 7.4, 150 mM NaCl) in solution in the absence of SLBs was incubated with TEV protease for 20 min. Protein mobility within the assembly after release from bilayer was measured by fluorescence recovery after photobleaching (FRAP). After 10 s of baseline recording (pre-bleach), a small area of interest (AOI) was photobleached. The average normalized fluorescence intensity of the AOI was plotted over time. (**B**) MBP-PrP-GFP was tethered via its C-terminal His-tag to SLBs. After washing to remove unbound MBP-PrP-GFP, TEV protease was added for 20 min. Cleavage of MBP-PrP-GFP was analyzed by Western blotting using an antibody against PrP. **(C, D)** MBP-PrP-GFP was anchored via its C-terminal His-tag to the NTA-Ni lipids in the SLBs and the MBP released with TEV protease. 3C protease was then added to cleave the C-terminal His-tag and to liberate PrP-GFP from the SLBs. The samples were analyzed by laser scanning microscopy after 3C protease treatment. Protein mobility within the assembly after release from the SLBs was measured by fluorescence recovery after photobleaching (FRAP). After 10 s of baseline recording (pre-bleach), a small area of interest (AOI) was photobleached. The average normalized fluorescence intensity of the AOI was plotted over time **(C)**. Volumetric 3D reconstructions generated with the IMARIS software. Red: SLBs, green: PrP-GFP **(D). (E)** Membrane anchored MBP-PrP-GFP (300 nM in 10 mM Tris pH 7.4, 150 mM NaCl) was incubated with 3C protease for 20 min. The samples were analyzed by laser scanning microscopy before and after incubation with 3C protease treatment (scale bar, 10 µm).

## Notes

### Competing Interest Statement

The authors have declared no competing interest.

## REFERENCES

Agarwal A, Rai SK, Avni A, Mukhopadhyay S (2021) An intrinsically disordered pathological prion variant Y145Stop converts into self-seeding amyloids via liquid-liquid phase separation. Proc Natl Acad Sci U S A 118

Alberti S, Dormann D (2019) Liquid-Liquid Phase Separation in Disease. Annual review of genetics 53: 171–194

Babinchak WM, Surewicz WK (2020) Liquid-Liquid Phase Separation and Its Mechanistic Role in Pathological Protein Aggregation. Journal of molecular biology 432: 1910–1925

Banik P, Ray K, Kamps J, Chen QY, Luesch H, Winklhofer KF, Tatzelt J (2024) VCP/p97 mediates nuclear targeting of non-ER-imported prion protein to maintain proteostasis. Life Sci Alliance 7

Banjade S, Rosen MK (2014) Phase transitions of multivalent proteins can promote clustering of membrane receptors. Elife 3

Barboni E, Rivero BP, George AJ, Martin SR, Renoup DV, Hounsell EF, Barber PC, Morris RJ (1995) The glycophosphatidylinositol anchor affects the conformation of Thy-1 protein. J Cell Sci 108 ( Pt 2): 487–497

Blochberger TC, Cooper C, Peretz D, Tatzelt J, Griffith OH, Baldwin MA, Prusiner SB (1997) Prion protein expression in Chinese hamster ovary cells using a glutamine synthetase selection and amplification system. Protein Eng 10: 1465–1473

Brandner S, Isenmann S, Raeber A, Fischer M, Sailer A, Kobayashi Y, Marino S, Weissmann C, Aguzzi A (1996) Normal host prion protein necessary for scrapie-induced neurotoxicity. Nature 379: 339–343

Brewis IA, Turner AJ, Hooper NM (1994) Activation of the glycosyl-phosphatidylinositol-anchored membrane dipeptidase upon release from pig kidney membranes by phospholipase C. Biochem J 303 ( Pt 2): 633–638

Case LB, Ditlev JA, Rosen MK (2019) Regulation of Transmembrane Signaling by Phase Separation. Annual review of biophysics 48: 465–494

Chesebro B, Trifilo M, Race R, Meade-White K, Teng C, LaCasse R, Raymond L, Favara C, Baron G, Priola S et al (2005) Anchorless prion protein results in infectious amyloid disease without clinical scrapie. Science 308: 1435–1439

Darling AL, Shorter J (2021) Combating deleterious phase transitions in neurodegenerative disease. Biochim Biophys Acta Mol Cell Res 1868: 118984

Ditlev JA (2021) Membrane-associated phase separation: organization and function emerge from a two-dimensional milieu. J Mol Cell Biol 13: 319–324

Durbin H, Young S, Stewart LM, Wrba F, Rowan AJ, Snary D, Bodmer WF (1994) An epitope on carcinoembryonic antigen defined by the clinically relevant antibody PR1A3. Proc Natl Acad Sci U S A 91: 4313–4317

Ferreira DG, Temido-Ferreira M, Miranda HV, Batalha VL, Coelho JE, Szego EM, Marques-Morgado I, Vaz SH, Rhee JS, Schmitz M et al (2017) alpha-synuclein interacts with PrPC to induce cognitive impairment through mGluR5 and NMDAR2B. Nature neuroscience

Furthmann N, Bader V, Angersbach L, Blusch A, Goel S, Sanchez-Vicente A, Krause LJ, Chaban SA, Grover P, Trinkaus VA et al (2023) NEMO reshapes the alpha-Synuclein aggregate interface and acts as an autophagy adapter by co-condensation with p62. Nat Commun 14: 8368

Gmachl M, Sagan S, Ketter S, Kreil G (1993) The human sperm protein PH-20 has hyaluronidase activity. FEBS Lett 336: 545–548

Hoyt F, Standke HG, Artikis E, Schwartz CL, Hansen B, Li K, Hughson AG, Manca M, Thomas OR, Raymond GJ et al (2022) Cryo-EM structure of anchorless RML prion reveals variations in shared motifs between distinct strains. Nat Commun 13: 4005

Huang JJ, Li XN, Liu WL, Yuan HY, Gao Y, Wang K, Tang B, Pang DW, Chen J, Liang Y (2020) Neutralizing Mutations Significantly Inhibit Amyloid Formation by Human Prion Protein and Decrease Its Cytotoxicity. Journal of molecular biology 432: 828–844

Jansen C, Parchi P, Capellari S, Vermeij AJ, Corrado P, Baas F, Strammiello R, van Gool WA, van Swieten JC, Rozemuller AJ (2010) Prion protein amyloidosis with divergent phenotype associated with two novel nonsense mutations in PRNP. Acta Neuropathol 119: 189–197

Kamps J, Lin YH, Oliva R, Bader V, Winter R, Winklhofer KF, Tatzelt J (2021) The N-terminal domain of the prion protein is required and sufficient for liquid-liquid phase separation: A crucial role of the Abeta-binding domain. J Biol Chem 297: 100860

Kocisko DA, Come JH, Priola SA, Chesebro B, Raymond GJ, Lansbury PT, Jr., Caughey B (1994) Cell-free formation of protease-resistant prion protein. Nature 370: 471–474

Konig AS, Rosener NS, Gremer L, Tusche M, Flender D, Reinartz E, Hoyer W, Neudecker P, Willbold D, Heise H (2021) Structural details of amyloid beta oligomers in complex with human prion protein as revealed by solid-state MAS NMR spectroscopy. J Biol Chem: 100499

Kostylev MA, Tuttle MD, Lee S, Klein LE, Takahashi H, Cox TO, Gunther EC, Zilm KW, Strittmatter SM (2018) Liquid and Hydrogel Phases of PrP(C) Linked to Conformation Shifts and Triggered by Alzheimer’s Amyloid-beta Oligomers. Mol Cell 72: 426–443 e412

Lauren J, Gimbel DA, Nygaard HB, Gilbert JW, Strittmatter SM (2009) Cellular prion protein mediates impairment of synaptic plasticity by amyloid-beta oligomers. Nature 457: 1128–1132

Lehto MT, Sharom FJ (1998) Release of the glycosylphosphatidylinositol-anchored enzyme ecto-5’-nucleotidase by phospholipase C: catalytic activation and modulation by the lipid bilayer. Biochem J 332 ( Pt 1): 101-109

Mallucci G, Dickinson A, Linehan J, Klohn PC, Brandner S, Collinge J (2003) Depleting neuronal PrP in prion infection prevents disease and reverses spongiosis. Science 302: 871–874

Matos CO, Passos YM, do Amaral MJ, Macedo B, Tempone MH, Bezerra OCL, Moraes MO, Almeida MS, Weber G, Missailidis S et al (2020) Liquid-liquid phase separation and fibrillation of the prion protein modulated by a high-affinity DNA aptamer. FASEB J 34: 365–385

Mead S, Lloyd S, Collinge J (2019) Genetic Factors in Mammalian Prion Diseases. Annual review of genetics 53: 117–147

Muller G, Bandlow W (1994) Lipolytic membrane release of two phosphatidylinositol-anchored cAMP receptor proteins in yeast alters their ligand-binding parameters. Arch Biochem Biophys 308: 504–514

Nygaard HB, Strittmatter SM (2009) Cellular prion protein mediates the toxicity of beta-amyloid oligomers: implications for Alzheimer disease. Arch Neurol 66: 1325–1328

Ondrejcak T, Klyubin I, Corbett GT, Fraser G, Hong W, Mably AJ, Gardener M, Hammersley J, Perkinton MS, Billinton A et al (2018) Cellular Prion Protein Mediates the Disruption of Hippocampal Synaptic Plasticity by Soluble Tau In Vivo. J Neurosci 38: 10595–10606

Passos YM, do Amaral MJ, Ferreira NC, Macedo B, Chaves JAP, de Oliveira VE, Gomes MPB, Silva JL, Cordeiro Y (2021) The interplay between a GC-rich oligonucleotide and copper ions on prion protein conformational and phase transitions. Int J Biol Macromol 173: 34–43

Polido SA, Stuani C, Voigt A, Banik P, Kamps J, Bader V, Grover P, Krause LJ, Zerr I, Matschke J et al (2024) Cross-seeding by prion protein inactivates TDP-43. Brain 147: 240–254

Prusiner SB (1998) Prions. Proc Natl Acad Sci U S A 95: 13363–13383

Purro SA, Nicoll AJ, Collinge J (2018) Prion Protein as a Toxic Acceptor of Amyloid-beta Oligomers. Biol Psychiatry 83: 358–368

Rambold AS, Miesbauer M, Rapaport D, Bartke T, Baier M, Winklhofer KF, Tatzelt J (2006) Association of Bcl-2 with misfolded prion protein is linked to the toxic potential of cytosolic PrP. Mol Biol Cell 17: 3356–3368

Rambold AS, Müller V, Ron U, Ben-Tal N, Winklhofer KF, Tatzelt J (2008) Stress-protective activity of prion protein is corrupted by scrapie-prions. EMBO J 27: 1974–1984

Ramos S, Kamps J, Pezzotti S, Winklhofer KF, Tatzelt J, Havenith M (2023) Hydration makes a difference! How to tune protein complexes between liquid-liquid and liquid-solid phase separation. Phys Chem Chem Phys 25: 28063–28069

Resenberger UK, Harmeier A, Woerner AC, Goodman JL, Muller V, Krishnan R, Vabulas RM, Kretzschmar HA, Lindquist S, Hartl FU et al (2011) The cellular prion protein mediates neurotoxic signalling of beta-sheet-rich conformers independent of prion replication. EMBO J 30: 2057–2070

Rogers M, Yehiely F, Scott M, Prusiner SB (1993) Conversion of truncated and elongated prion proteins into the scrapie isoform in cultured cells. Proc Natl Acad Sci U S A 90: 3182–3186

Smith LM, Strittmatter SM (2017) Binding Sites for Amyloid-beta Oligomers and Synaptic Toxicity. Cold Spring Harb Perspect Med 7

Snead WT, Gladfelter AS (2019) The Control Centers of Biomolecular Phase Separation: How Membrane Surfaces, PTMs, and Active Processes Regulate Condensation. Mol Cell 76: 295–305

Stohr J, Watts JC, Legname G, Oehler A, Lemus A, Nguyen HO, Sussman J, Wille H, DeArmond SJ, Prusiner SB et al (2011) Spontaneous generation of anchorless prions in transgenic mice. Proc Natl Acad Sci U S A 108: 21223–21228

Tange H, Ishibashi D, Nakagaki T, Taguchi Y, Kamatari YO, Ozawa H, Nishida N (2021) Liquid-liquid phase separation of full-length prion protein initiates conformational conversion in vitro. J Biol Chem: 100367

Tatzelt J, Prusiner SB, Welch WJ (1996) Chemical chaperones interfere with the formation of scrapie prion protein. EMBO J 15: 6363–6373

Walmsley AR, Zeng FN, Hooper NM (2001) Membrane topology influences N-glycosylation of the prion protein. Embo J 20: 703–712

Wang X, Jansen G, Fan J, Kohler WJ, Ross JF, Schornagel J, Ratnam M (1996) Variant GPI structure in relation to membrane-associated functions of a murine folate receptor. Biochemistry 35: 16305–16312

Winklhofer KF, Heske J, Heller U, Reintjes A, Muranji W, Moarefi I, Tatzelt J (2003) Determinants of the in vivo-folding of the prion protein: a bipartite function of helix 1 in folding and aggregation. J Biol Chem 278: 14961–14970

Zbinden A, Perez-Berlanga M, De Rossi P, Polymenidou M (2020) Phase Separation and Neurodegenerative Diseases: A Disturbance in the Force. Dev Cell 55: 45–68

